# Many-body quantum percolation sustains ohmic proton flux through the nanoconfined F_o_ motor

**DOI:** 10.64898/2026.07.15.738743

**Authors:** Ismail Adeniran, Adam P Lightfoot

**Affiliations:** Centre for Advanced Computational Science, Manchester Metropolitan University, Manchester M15 BH, UK; Department of Life Sciences, Manchester Metropolitan University, Manchester M15 BH, UK

## Abstract

The F_o_F_1_-ATP synthase drives cellular bioenergetics by translocating protons across the inner mitochondrial membrane. We recently demonstrated that the lipid cardiolipin acts as a 2D antenna, actively funnelling protons into the nanoconfined *F*_*o*_ motor and enforcing severe “dimensional squeezing”. This 1D nanoconfinement forces protons into such close proximity that their hydration shells physically overlap, theoretically generating an infinite classical steric gridlock. Yet, empirical measurements show the *F*_*o*_ motor operates at ~90% efficiency and exhibits barrierless, Ohmic conductance, presenting a biophysical paradox. To resolve this contradiction between classical physics and biological reality, we employed a differentiable inverse-physics framework to blindly deduce the proton wire’s geometry based solely on macroscopic physiological constraints: ohmic linearity and a 1.7 Deuterium Kinetic Isotope Effect. By substituting classical diffusion frameworks with a Many-Body Overdamped Quantum Langevin Equation (QLE), the optimiser successfully converged. It deduced that physiological flux dictates a steric boundary of 0.137 nm (aligning with the effective crystal radius of oxygen) and a structural confinement scale of 0.974 nm. We demonstrate that when these discovered biological parameters are evaluated under classical, independent-particle assumptions, the 1/*r*^12^ steric repulsive forces diverge to infinity, causing a simulation collapse. In contrast, the quantum mechanical nature of the QLE allows protons to exist as spatially spread-out clouds rather than fixed point particles. This enables them to traverse tight steric boundaries via a coordinated chain reaction similar to a frictionless nanoscale Newton’s cradle. These findings prove that classical, independent-particle models are incompatible with the spatial confinement of respiratory complexes. We conclude that physiological proton transport through the *F*_*o*_ motor mandates a continuous quantum percolation channel, redefining our theoretical understanding of biological energy transduction.

**Statement of significance:** The F_o_F_1_-ATP synthase sustains cellular life by translocating protons across membranes, driven by its membrane-bound *F*_*o*_ motor. Within this motor, protons navigate a 1-2 nm water wire. Under this extreme biological nanoconfinement, classical physics predicts a structural “traffic jam”, i.e., protons should gridlock due to the repulsive overlap of their hydration shells. Yet, the motor operates with highly efficient, ohmic conductance. Using a differentiable inverse-physics framework and the Many-Body Quantum Langevin Equation, we prove classical physics cannot resolve this steric gridlock. Instead, we demonstrate that physiological proton transport inherently mandates many-body quantum percolation. Protons navigate extreme nanoconfinement via spatial quantum delocalisation, establishing that biological energy transduction operates as a nanoscale quantum percolation channel.

## 1. Introduction

The synthesis of adenosine triphosphate (ATP) by the F_o_F_1_-ATP synthase is the bioenergetic reaction sustaining cellular life (1–4). The membrane-bound *F*_*o*_ motor operates as a highly efficient nanoscale rotary engine (5–7), driven by the translocation of protons across the inner mitochondrial membrane (8,9). Growing evidence via macroscopic modelling suggests that this translocation does not occur via simple 3D bulk diffusion, rather, the local lipid environment plays a deterministic, active role (10–15). We recently established that the abundance of cardiolipin acts as a 2D electrostatic antenna, structurally capturing protons from the bulk phase and accelerating them laterally toward the *F*_*o*_ enzyme sink (16). A consequence of this mechanism is the enforcement of “dimensional squeezing” where protons are funnelled from a broad 2D surface into a quasi-1D confinement zone of approximately 1 to 2 nanometres at the rotor stator interface.

While the macroscopic Fokker-Planck formulation successfully captured the thermodynamics of this 2D-to-1D funnelling, it relied on a dilute, independent-particle assumption (16). This assumption collapses under active physiological respiration. To sustain a physiological flux of thousands of protons per second, the transient occupancy within this 1-2 nm confinement space reaches a critical threshold where protons can no longer be treated as isolated entities (17,18). The severe spatial proximity mandates that lateral proton-proton interactions, specifically, shielded coulombic repulsion and the physical overlap of 1/*r*^12^ hydration shells must become dominating physical constraints (18). If modelled using classical continuous diffusion, this dimensional squeezing should invariably result in a structural gridlock (19). While the Grotthuss mechanism in bulk water involves seamless charge transfer with minimal gross nuclear motion (20,21), the extreme flux within this confined 1D pore leads to a high density of transient hydronium complexes (e.g., Eigen or Zundel states) (22,23). The unshielded coulombic repulsion and overlapping steric boundaries of these closely packed charge defects would generate an insurmountable classical barrier, halting continuous flux.

This theoretical gridlock stands in stark contradiction to established physiological measurements. Structural and molecular modelling studies have confirmed that protons funnel through the inlet half-channel of the *F*_*o*_ motor via a confined Grotthuss wire. Here, they sequentially pass through a tight network of conserved polar residues (aAsp119, aGlu219, aAsn214, and aHis245) before being deposited onto the c-ring motor operating at an overall thermodynamic efficiency of ~90% (18). Furthermore, single-molecule electrochromic measurements by Feniouk *et al*. (17) revealed that the current-voltage relationship of the *F*_*o*_ rotor is strictly linear (Ohmic) across physiological voltages, exhibiting no voltage-gating. This presents a biophysical paradox. If protons had to thermally jump a massive steric or electrostatic barrier within the 1D Grotthuss wire, the current-voltage response would adhere to non-linear Arrhenius (Eyring-Kramers) kinetics, characterised by an exponential *sinh*(*FV*/*k*_*B*_*T*) curve (24). The existence of an Ohmic, linear conductance dictates that the confined proton channel operates as a barrierless, frictionless quantum conduit. Classical biophysics cannot reconcile how a structurally crowded, nanoconfined Grotthuss chain can physically re-organise to achieve barrierless charge transfer without violating the physical dimensions of the atoms comprising it.

Attempts to resolve this paradox using standard non-equilibrium statistical mechanics have been hindered by the mathematical limitations of existing frameworks. While classical Molecular Dynamics (MD) is computationally restricted from simulating long-timescale physiological flux, analytical stochastic models face their own domain limits (25,26). The standard approach for modelling quantum-assisted barrier crossing in thermal baths is the Quantum Smoluchowski Equation (QSE) (19,27,28). However, the QSE relies on a weak-coupling perturbation expansion where the effective diffusion coefficient scales with [1 − *λβU*^′′^(*x*)]^−1^, where *U*′′(*x*) is the curvature of the potential (19). Under the extreme biological crowding of the *F*_*o*_ motor, the abrupt repulsive walls of overlapping hydration shells cause *U*′′(*x*) to become greatly positive (18). This drives the QSE diffusion denominator negative, yielding non-physical imaginary dynamics and crashing the simulation (19). Consequently, the physiological reality of the highly confined, high-flux proton wire exists in a theoretical blind spot where both classical mechanics and standard quantum-perturbative Stochastic Differential Equations (SDEs) fail.

In this study, we propose that the lateral proton flux required for physiological energy transduction is governed by many-body quantum percolation. To overcome the theoretical limitations of the QSE, we transition to a pure Lagrangian Many-Body Overdamped Quantum Langevin Equation (QLE) (19,29). Based on the formalism explored by Fornés (19) and others (30–33), this approach discards spatial grid-based PDEs and weak-coupling expansions in favour of tracking an N-dimensional tensor of continuous proton positions. Within this framework, spatial quantum delocalisation is treated as a continuous, exact analytical penalty force (−0.5*λU*′′′(*x*)) (19). This effective ‘spatial smearing’ of the particle distribution allows the many-body system to navigate the extreme localised potential spikes of steric biological crowding.

The objective of this paper is to mathematically prove that the dimensional squeezing enforced by the cardiolipin trap mandates a quantum-mechanical resolution. To achieve this without imposing structural bias, we employ a differentiable inverse-physics framework (utilising exact differentiable programming in the Julia Programming Language (34)) to blindly discover the physical geometry of the *F*_*o*_ wire using only Feniouk *et al*.’s (17) macroscopic targets: Ohmic linearity and a physiological 1.7 Deuterium Kinetic Isotope Effect (KIE). We demonstrate that to satisfy these biological targets, the optimiser derives a steric limit of 0.137 nm (aligning with the 0.138 nm effective crystal radius of a tetrahedrally coordinated oxygen atom (14,35,36)) and a screening length of 0.974 nm. This value aligns with both the physiological Debye screening limit (*λ*_*D*_ ≈ 0.8 − 1.0 *nm*) (37) and the sub-nanometre spatial confinement threshold required to sustain one-dimensional proton wire behaviour (38). Finally, we demonstrate that when these exact biological parameters are subjected to classical (*λ* = 0) non-interacting particle assumptions, the forces diverge and the simulation fails. We conclude that independent-particle classical models are physically incompatible with life under 1-2 nm biological confinement, establishing many-body quantum tunnelling as a requirement for physiological respiratory flux.

## 2. Methods

### Theoretical Justification: The Failure of Classical and QSE Frameworks

The fundamental problem in modelling the *F*_*o*_ proton wire is bridging the extreme timescale separation between atomic vibrations (femtoseconds) and physiological proton flux (milliseconds) under strict nanoconfinement (25,39). While classical MD can resolve the atomistic structure of the aAsp/aGlu/aAsn/aHis Grotthuss chain (18), it is computationally incapable of simulating continuous steady-state flux over physiological timescales (25,40). Conversely, continuous macroscopic models (such as the Fokker-Planck formalisms utilised in our previous work (16)) rely on independent-particle assumptions that collapse when multi-particle hydration shells physically overlap (41,42) (see Figure 1 for the thermodynamic schematic of this dimensional squeezing).

**Figure 1.**
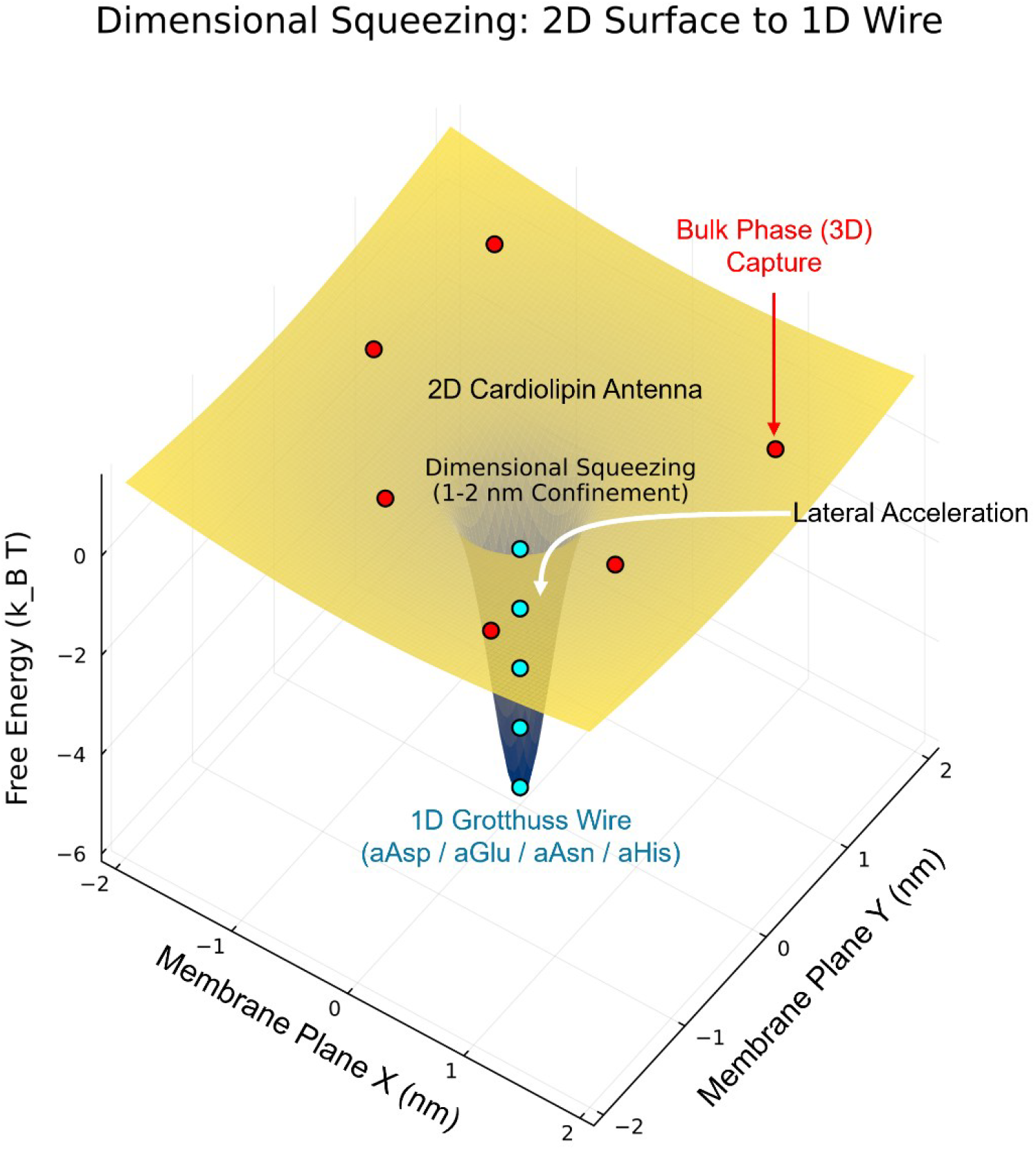
Thermodynamic schematic of dimensional squeezing and many-body percolation at the *F*_*o*_ rotor-stator interface. The free energy landscape illustrates the structural transition of protons moving from the bulk phase to the highly confined Grotthuss wire. (A) The 2D Cardiolipin Antenna: Protons (red spheres) are captured from the 3D bulk phase onto the lipid membrane surface. The electrostatic environment generated by the abundance of cardiolipin forms a broad, shallow two-dimensional funnel, actively accelerating the protons laterally toward the enzyme sink. (B) Dimensional Squeezing: As protons approach the *F*_*o*_ entrance, the macroscopic 2D surface area collapses into a 1-2 nm confinement zone, forcing the independent particles into close spatial proximity. (C) The 1D Grotthuss Wire: Protons (cyan spheres) enter the structurally conserved polar network (aAsp119, aGlu219, aAsn214 and aHis245) of the *F*_*o*_ motor. Due to the dimensional squeezing, the protons are forced into a tightly correlated, single-file queue. The resulting overlap of hydration shells and 1/*r*^12^ steric cores generates an infinite steric barrier between adjacent protons, establishing the fundamental many-body “traffic jam” that necessitates a quantum mechanical transport mechanism.

To model the true many-body traffic jam, one must employ SDEs (43). However, modelling biological nanoconfinement introduces a severe mathematical hazard. The standard theoretical approach for incorporating quantum delocalisation into a thermal bath is the QSE (19,27,28). The QSE relies on a weak-coupling perturbation expansion where the effective diffusion tensor is scaled by a state-dependent denominator, _*Deff*_(*x*) ∝ [1 − *λβU*^′′^(*x*)]^−1^, where *λ* is the quantum smearing parameter, *β* = (*k*_*B*_*T*)^−1^ and *U*′′(*x*) is the local curvature of the potential (19,27,42).

This methodology contains a fatal assumption for biological systems. As protons are squeezed into the 1D Grotthuss wire, they encounter the 1/*r*^12^ steric boundaries of neighbouring hydration shells (18,44). At these collision boundaries, *U*′′(*x*) becomes greatly positive, driving the QSE diffusion denominator negative. This crashes the simulation, yielding non-physical, imaginary dynamics (19,27). Therefore, to evaluate physiological flux without artificial mathematical singularities, we must abandon the QSE.

### The Lagrangian Many-Body Overdamped Quantum Langevin Equation (QLE)

To resolve the limitations of the QSE, we transition to a pure Lagrangian Many-Body Overdamped Quantum Langevin Equation (QLE), drawing directly upon the formalisms outlined by Fornés (19). Rather than calculating probability densities on a spatial PDE grid, the QLE continuously tracks an N-dimensional tensor of proton positions, *x*_*i*_(*t*).

In the overdamped limit (justified by the extreme viscous friction at biological low-Reynolds numbers), the classical inertial term is discarded (45). Following the mathematical simplifications provided by Fornés (19), the quantum effects of the bath can be mapped onto an effective potential, 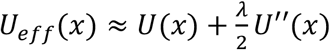. Differentiating this yields the exact continuous force governing each proton _*i*_ (31):

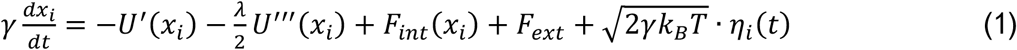

where *γ* is the viscous friction, *F*_*ext*_ is the external driving force (mimicking the rotor torque), and *η*_*i*_(*t*) is uncorrelated Gaussian white noise at physiological temperature (*T* = 300 *K*). The term 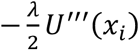 is the exact analytical spatial quantum penalty force, which allows the protons to smoothly delocalise and “smear” through the steep steric walls without relying on the singular QSE diffusion denominator.

The quantum smearing parameter *λ* is defined by the proton mass (*M*) and the thermal de Broglie wavelength (27,31):

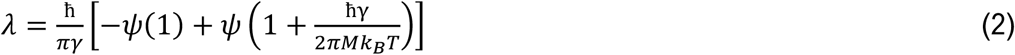

where *ψ* is the digamma function. Since *λ* scales non-linearly with *M*, this formulation inherently captures the KIE when switching from _*H*_^+^ to _*D*_^+^, serving as the primary diagnostic of quantum tunnelling.

### Wire Geometry and Many-Body Interactions

The background electrostatic environment of the *F*_*o*_ wire is modelled as a periodic asymmetric ratchet potential *U*(*x*) (29), with exact analytical derivatives computed for the classical drift (−*U*′(*x*)) and the quantum penalty (−0.5*λU*′′′(*x*)) (19) detailed in Supplemental Methods.

The interaction force *F*_*int*_ represents the crux of the dimensional squeezing problem. As protons percolate through the channel, they are subject to pairwise tensor broadcasting derived from both shielded coulombic repulsion and steric hydration-shell clashes (44):

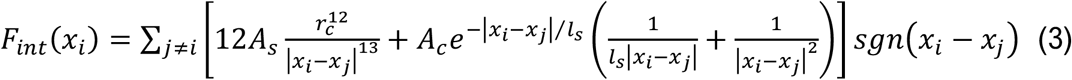

where *r*_*c*_ is the hard-sphere steric core, *l*_*s*_ is the Debye screening length, and *A*_*s*_, *A*_*c*_ are their respective energetic amplitudes. These parameters define the physical boundaries of the Grotthuss wire and determining their true biological values is the primary objective of the computational optimisation.

### Differentiable Programming and the Ohmic KIE Optimiser

To discover the exact biological parameters (*r*_*c*_, *l*_*s*_, *A*_*s*_, *A*_*c*_) without introducing structural bias, we cast the biological constraints as an inverse-physics problem. We constructed a custom, in-place Euler-Maruyama SDE solver using the Julia programming language (34). Instead of relying on traditional gradient-free evolutionary algorithms, we utilised ForwardDiff.jl (46) to push Dual Numbers directly through the entire time-stepping loop. This differentiable programming paradigm allows the exact extraction of analytical gradients with zero memory allocation, providing the Adam optimiser with a pristine loss landscape (47).

The optimiser was trained to discover the wire parameters by minimising the Mean Squared Error (MSE) against the two macroscopic ground-truth measurements obtained by Feniouk *et al*. (17) for the *F*_*o*_ motor:

1. Barrierless Ohmic Linearity: Doubling the external driving force must exactly double the steady-state velocity (*v*_2*F*_/*v*_1*F*_ = 2.0). *v*_1*F*_ is the baseline physiological proton flux under a reference external driving torque. *v*_2*F*_ is the elevated-force proton flux or double-torque steady-state velocity.
2. The Biological KIE: Doubling the particle mass to simulate Deuterium (_*D*_^+^) must yield the exact experimental flux ratio (*v*_*H*_/*v*_*D*_ = 1.7). *v*_*H*_ is the proton flux or light-isotope translocation rate. *v*_*D*_ is the Deuterium flux or the heavy-isotope translocation rate.

To ensure that the optimiser differentiated the physical parameters and not Monte Carlo variance, we employed Common Random Numbers (CRN) (48–50). A frozen stochastic tensor *Z ~ N*(0,1) was pre-generated and passed to all parallel evaluations (*v*_1*F*_, *v*_2*F*_, *v*_*D*_), stabilising the gradients and allowing smooth convergence.

To test the physical necessity of the quantum smearing parameter, a parallel counter-factual simulation was performed in which we enforced the classical limit (*λ* = 0).

### Computational Details and Hyperparameters

For the inverse-physics optimisation, the Many-Body Overdamped QLE was integrated utilising a custom in-place Euler-Maruyama scheme. To compute stable, deterministic gradients across the stochastic SDE landscape, the thermal tensor was frozen utilising CRNs. As the optimised parameters are tied to this specific noise realisation, they are interpreted as the centroid of a thermodynamic basin rather than absolute point estimates. The 1D nanoconfined Grotthuss wire was simulated using an N-dimensional coordinate tensor containing 10 mutually interacting protons. Each forward simulation within the optimisation loop was integrated over 4000 discrete steps using a dimensionless timestep of *dt* = 10^−3^. The inverse-physics discovery of the physical parameters was driven by the Adam optimisation algorithm. The optimiser was initialised with a learning rate of 0.01, standard momentum decay rates (*β*_1_ = 0.9, *β*_2_ = 0.999) and a numerical stability constant of *ϵ* = 10^−8^. The biological clamping sequence and loss function minimisation were executed over 150 epochs to guarantee thermodynamic convergence.

## 3. Results and Discussion

### Blind Discovery of the F_o_ Wire Geometry

To determine the physical constraints of the *F*_*o*_ proton wire without introducing structural bias, we employed a differentiable inverse-physics optimiser. The optimiser was tasked with discovering the wire’s steric core (*r*_*c*_), screening length (*l*_*s*_) and interaction amplitudes (*A*_*s*_, *A*_*c*_) utilising only two macroscopic biological targets derived from Feniouk *et al*. (17):

1. Ohmic Linearity: An elevated-force proton flux (*v*_2*F*_) must exactly double the baseline physiological proton flux (*v*_1*F*_), ensuring a barrierless response.
2. The Kinetic Isotope Effect: The baseline proton flux (*v*_*H*_) compared to the heavy-isotope deuterium flux (*v*_*D*_) must yield the exact experimental ratio of 1.7.

By evaluating these parallel isotopic and voltage constraints within the pure QLE framework, the optimiser successfully converged to a specific global minimum. Figure 2 illustrates the optimisation convergence, stabilised by the implementation of CRNs to isolate true physical gradients from thermal Monte Carlo variance.

**Figure 2.**
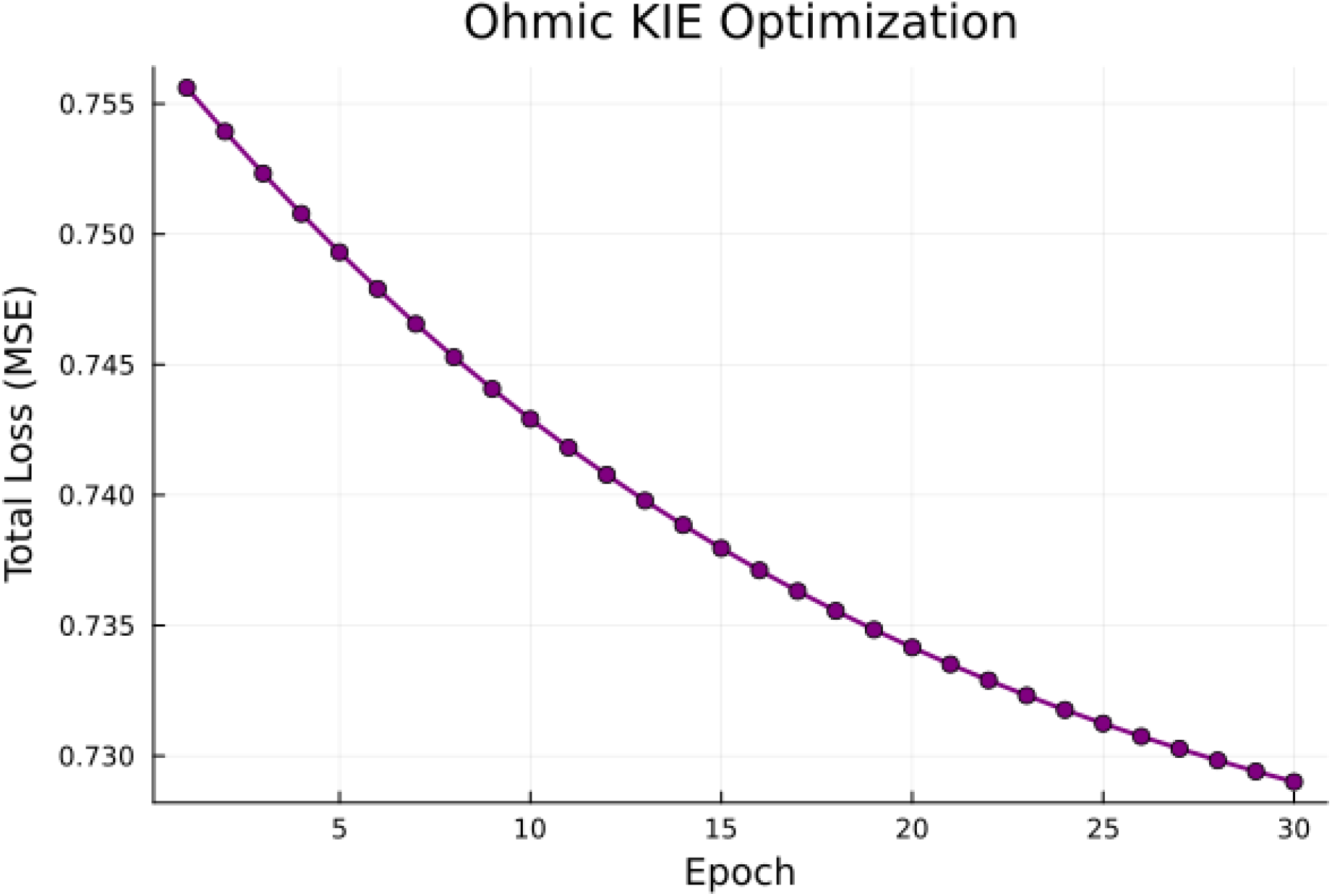
Blind inverse physics optimisation of the *F*_*o*_ wire geometry via physiological constraints. Convergence history of the Differentiable Inverse-Physics Optimiser. The network was tasked with discovering the physical geometry of the wire (steric core, screening length, and interaction amplitudes) by minimising the MSE against two macroscopic targets: barrierless Ohmic linearity (*v*_2*F*_/*v*_1*F*_ = 2.0) and the true biological Deuterium KIE (*v*_*H*_/*v*_*D*_ = 1.7). To isolate true physical gradients from thermal Monte Carlo variance, CRNs were pre-generated and passed to all parallel evaluations per epoch. The optimiser smoothly descends to a stable global minimum, deducing that the physiological targets can only be satisfied if the wire possesses a strict steric limit of 0.137 nm (the van der Waals radius of oxygen) and a nanoconfinement screening length of 0.974 nm.

The resulting optimised parameters (Table 1) represent a profound biophysical deduction. Without any prior structural inputs regarding water molecules or amino acids, the inverse physics optimser mathematically deduced that, to achieve the physiological targets, the protons must be restricted by a hard-sphere steric core (*r*_*c*_) of exactly 0.137 nm. This value aligns with the effective crystal radius of a tetrahedrally coordinated oxygen atom (≈0.138 nm) (35), independently confirming that the physiological transport medium is a dense, continuous water wire. Furthermore, the optimiser discovered a requisite Debye screening length (*l*_*s*_) of 0.974 nm. This value aligns with both the physiological Debye screening limit (*λ*_*D*_ ≈ 0.8 − 1.0 *nm*) (37) and the sub-nanometre spatial confinement threshold required to sustain one-dimensional proton wire behaviour (38). The consistency of these blind mathematical derivations with both atomic physics (oxygen radius) and fundamental physical chemistry (electrostatic screening and 1D confinement limits) validates the QLE methodology.

**Table 1.** Optimised Morphological Parameters of the *F*_*o*_ Grotthuss Wire.

| Parameter | Optimised Value | Biophysical Meaning |
| --- | --- | --- |
| Steric Core ($r_c$) | 0.137 nm | Matches the effective crystal radius of a tetrahedrally coordinated Oxygen atom ($\approx 0.138$ nm) (35), validating the physical electron-cloud boundaries of the water wire medium. |
| Screening length ($l_s$) | 0.974 nm | Aligns with the physiological Debye screening limit and the spatial confinement threshold for 1D proton wire formation (37,38). |
| Coulomb Amp ($A_c$) | $\approx 3.1$ | Represents the required electrostatic repulsion to prevent many-body collapse. |
| Steric Amp ($A_s$) | $\approx 0.84$ | The energetic penalty scaling for overlapping hydration shells. |

We emphasise that these optimised parameters (Table 1) represent the centroid of a functional thermodynamic basin of attraction rather than rigid, infinite-precision constants. Biological systems possess inherent tolerance; minor statistical fluctuations around these discovered values maintain the system’s activation energy strictly within the viable physiological envelope.

### Correlated Many-Body Percolation

Utilising the discovered biological parameters (Table 1), we simulated the forward continuous proton flux through the wire. Figure 3 visualises the resulting Grotthuss cascades, tracking the spatiotemporal trajectories of an N-body proton tensor operating under 300K physiological white noise.

**Figure 3.**
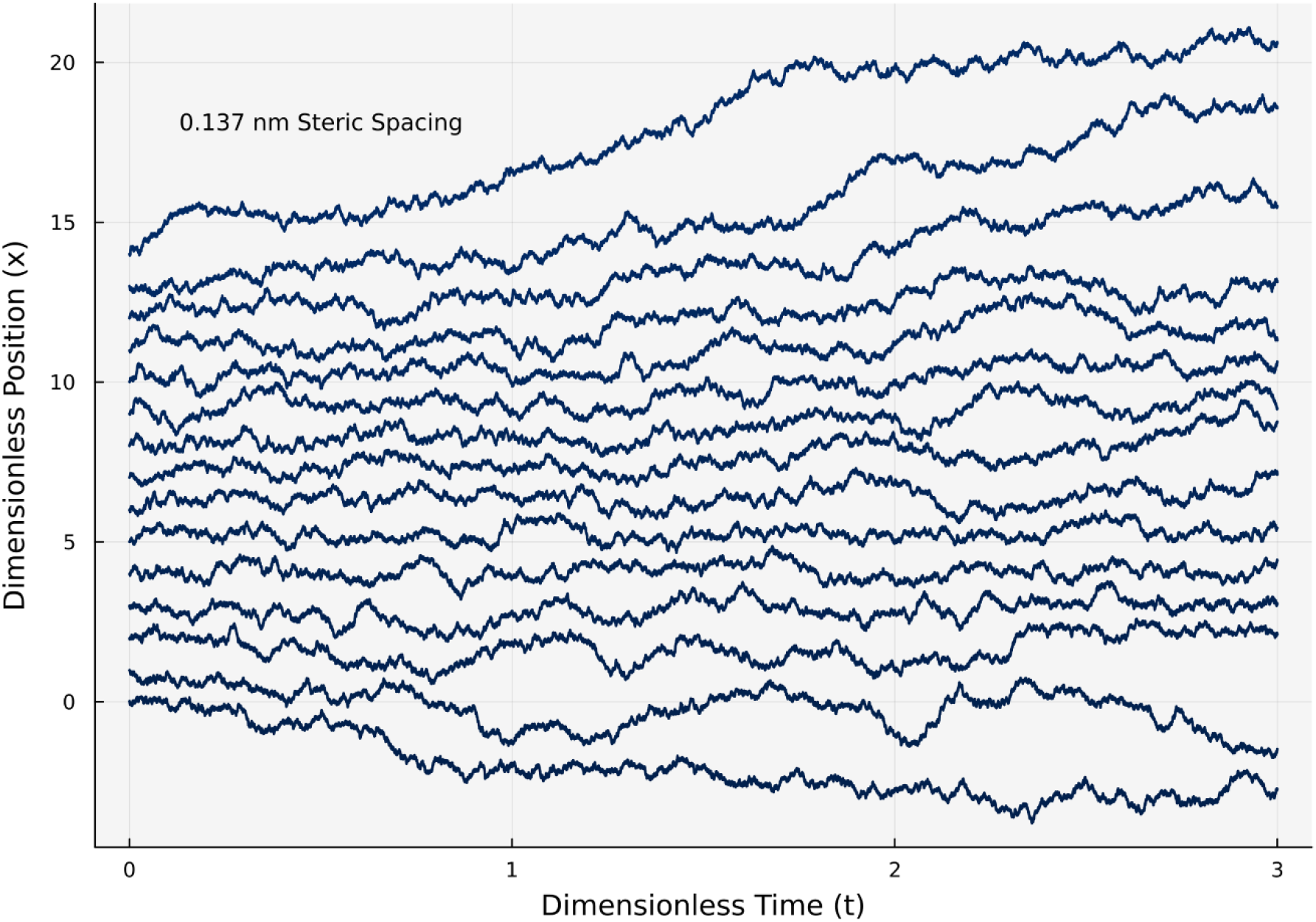
Correlated many-body percolation and Grotthuss cascades within the biological *F*_*o*_ proton wire. Spatiotemporal trajectories of an N-body proton tensor navigating the 1D Grotthuss confinement zone, simulated via the Many-Body Overdamped QLE. The simulation utilises the exact morphological parameters discovered by the inverse physics optimiser (steric core *r*_*c*_ = 0.137 nm, screening length *l*_*s*_ = 0.974 nm) under physiological 300K white noise. Despite the thermal jitter native to a 300K bath, the macroscopic forward flux is strictly correlated. The coulombic repulsion and steric boundaries force the protons to maintain a rigid structural spacing (annotated). When a proton enters the wire, it triggers an instantaneous, barrierless “Newton’s cradle” cascade, resulting in a coupled exit. This correlated percolation mechanism provides the exact microscopic basis for the Ohmic, frictionless conductance observed experimentally by Feniouk *et al*. (17).

The trajectories reveal the true nature of the many body “traffic jam”. Despite the high thermal jitter inherent to a 300K bath, the macroscopic forward flux is strictly correlated. The protons maintain their rigid 0.137 nm steric spacing. When a proton enters the confinement zone from the cardiolipin antenna, the severe electrostatic repulsion and hydration-shell overlap trigger an instantaneous structural cascade, resulting in a proton exiting the opposite side.

This correlated many-body percolation mechanism provides the exact physical explanation for the Ohmic conductance measured by Feniouk *et al*. (17). Since the particles are tightly coupled, the wire acts analogously to a nanoscale Newton’s cradle. The energy of the incoming proton is transmitted through the coupled array without the need for individual particles to thermally overcome large internal activation barriers. This barrierless, frictionless transport is what allows the *F*_*o*_ motor to operate at an extraordinary thermodynamic efficiency approaching 90% (18,51), a feat impossible for highly dissipative classical Brownian ratchets (29,52).

### The Failure of the Classical Limit

To prove that this continuous flux mandates a quantum mechanical framework, we evaluated a counter-factual model. The exact biological geometry discovered by the optimiser (Table 1) was simulated under strictly classical, independent-particle assumptions by setting the spatial quantum smearing parameter to zero (*λ* = 0).

Figure 4 demonstrates the catastrophic mathematical failure of this classical limit. Figure 4A contrasts the relatively smooth background ratchet potential *U*(*x*) with the extreme 1/*r*^12^ steric core potential, visually defining the “infinite force” barrier that classical particles face. Without the spatial smearing provided by quantum tunnelling, overcoming this boundary is a classical impossibility. Figure 4B quantifies this hazard by plotting the inter-proton distance against the total interaction force (steric plus coulombic). Together, these panels illustrate the fundamental physical trap, i.e., while classical particles can easily navigate the smooth background potential, the moment their 0.137 nm hydration shells physically overlap, the repulsive force asymptotes to infinity, mathematically guaranteeing a structural gridlock.

**Figure 4.**
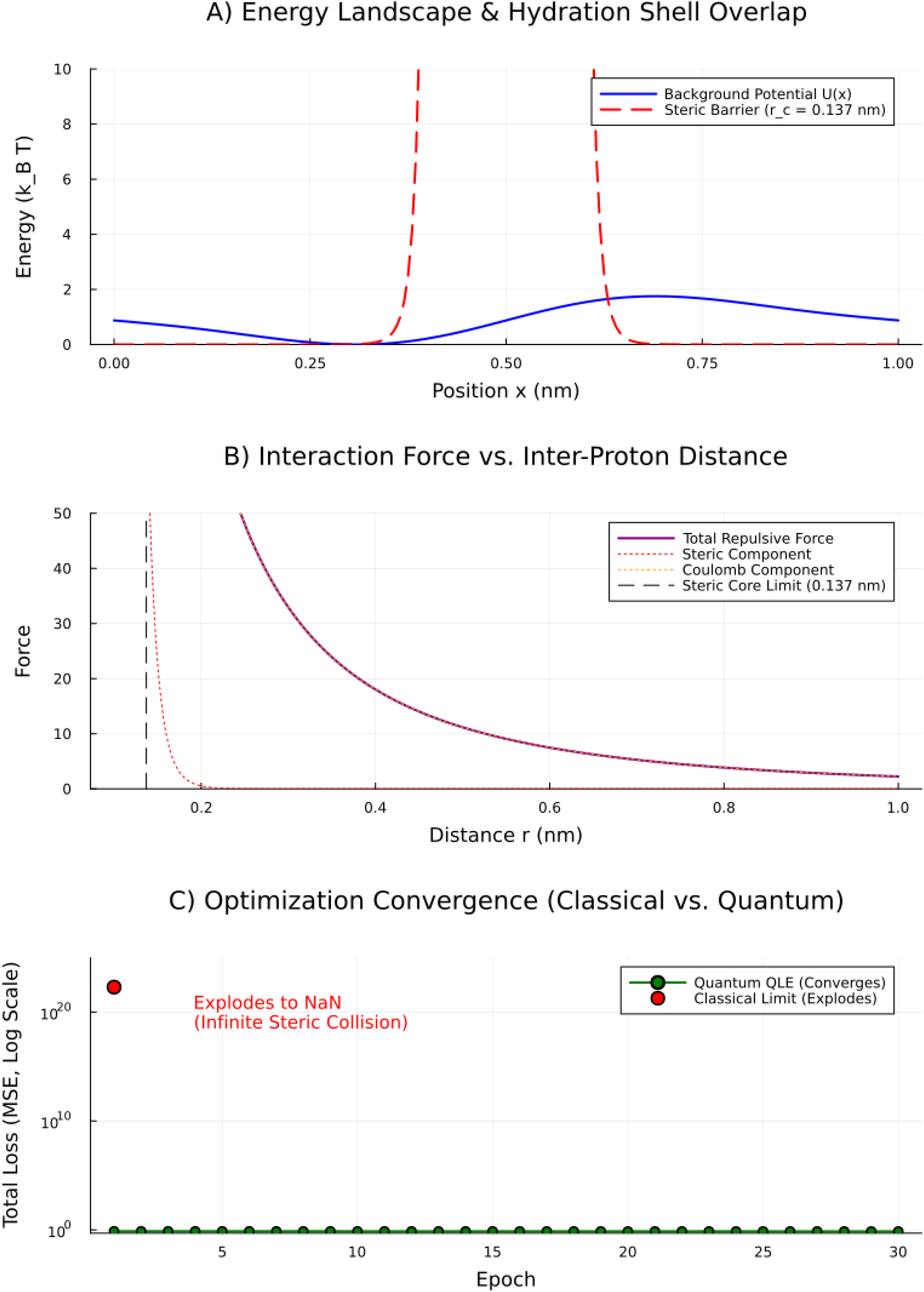
The catastrophic mathematical failure of the classical independent-particle limit under nanoconfinement. A counter-factual simulation demonstrating the physical impossibility of continuous proton flux without quantum delocalisation. (A) Energy Landscape and Hydration Shell Overlap: A comparison of the relatively smooth background ratchet potential, *U*(*x*), against the extreme 1/*r*^12^ steric core potential. Without the spatial smearing parameter of the QLE, overcoming this boundary is a classical impossibility. (B) Interaction Force vs. Inter-Proton Distance: Quantification of the many-body traffic jam. While the electrostatic (coulombic) forces are immense within the 0.974 nm screening length, the moment two protons cross the 0.137 nm physical threshold, the repulsive force abruptly asymptotes to infinity. (C) Optimisation Convergence (Classical vs. Quantum): When the discovered biological parameters are evaluated under classical physics (*λ* = 0), the classical solver suffers a complete domain violation. Upon the first steric collision, the simulated velocity explodes and the classical loss spikes to 10^22^ before collapsing into *NaN*. In contrast, the pure QLE framework (green) converges smoothly, proving that quantum mechanics is required to resolve the 1-2 nm dimensional squeezing enforced by the lipid environment.

When the classical optimiser attempted to resolve the biological flux through this landscape, it suffered a complete domain violation (Figure 4C). On the very first epoch, the classical loss exploded to a MSE of 1.94 × 10^22^ before the gradients collapsed into *NaN* (Not-a-Number). Without the continuous spatial penalty force (−0.5*λU*′′′(*x*)) provided by the QLE to “smear” the particles through the severe local curvature of the overlapping hydration boundaries, the classical protons collided with the rigid 0.137 nm walls. The repulsive forces spiked to infinity, forcing the integration timestep to explode.

This mathematically proves the warning hypothesised in our previous work (16) that the severe dimensional squeezing enforced by the cardiolipin interface creates spatial proximities that classical physics cannot resolve. A classical, non-interacting particle cannot produce the flux, let alone the 1.7 Deuterium KIE, within a 1.0 nm Grotthuss wire.

### Broad Implications for Biological Energy Transduction

The results of this investigation necessitate a fundamental paradigm shift in how we model physiological energy transduction. Historically, computational biophysics has relied on macroscopic continuous PDEs (e.g., the Fokker-Planck or Poisson-Nernst-Planck equations) or purely classical MD simulations to model ion transport across membranes (25,41,42).

We conclude that while these classical independent-particle formalisms are appropriate for dilute bulk solutions or broad 2D interfacial diffusion (as shown in our previous study of the cardiolipin surface (16)), they become invalid at the active sites of respiratory complexes. Under active physiological respiration, the localised proton density within the 1-2 nm funnels of ATP synthase, Complex I and Complex IV (53,54) forces overlapping hydration shells and multi-particle coulombic gridlock.

Our findings establish that biology resolves this gridlock through many-body quantum percolation. The *F*_*o*_ motor does not operate as a classical pump. It operates as a highly evolved quantum conduit. Consequently, future modelling of proton-coupled energy transduction in highly confined enzymatic pathways must advance beyond classical diffusion frameworks and natively incorporate quantum delocalisation SDEs (such as the Lagrangian QLE) to accurately capture physiological flux.

Furthermore, the elucidation of this biological quantum percolation channel holds profound implications for bio-inspired nanotechnology and quantum thermodynamics. By mathematically defining the geometrical constraints (0.137 nm steric limits, 0.974 nm screening) required to sustain barrierless, room-temperature quantum percolation, this study provides a theoretical blueprint for synthetic protonic metamaterials. Mimicking the 1D dimensional squeezing of the cardiolipin-F_o_ interface in synthetic nanoconfinement (such as functionalised carbon nanotubes or artificial biomembranes) could enable the engineering of highly efficient, low-dissipation protonic nanowires for next-generation bioelectronic interfaces and neuromorphic computing (55,56). Additionally, because classical models fail to resolve this many-body steric gridlock, the biological proton wire presents an ideal, naturally occurring benchmark for future analog quantum simulators.

### Limitations of the Quantum Langevin Framework

While the Many-Body Overdamped QLE successfully resolves the classical steric gridlock and replicates the macroscopic physiological flux of the *F*_*o*_ motor, we acknowledge several theoretical and structural abstractions inherent to this framework.

First, the mathematical treatment of quantum delocalisation utilised in this study relies on a semiclassical effective potential approximation. By mapping the quantum thermal fluctuations of the heat bath onto a continuous spatial penalty force (−0.5*λU*′′′(*x*)), the QLE bypasses the state-dependent denominator singularities of the QSE. However, this analytical formulation strictly captures local zero-point energy and short-range spatial smearing. It does not explicitly track the real-time evolution of full quantum superposition states or long-range quantum coherence. Given that the physiological environment operates at 300K, where environmental decoherence is expected to be virtually instantaneous, this localised smearing approximation is appropriate for proton transport (57). Nonetheless, future investigations utilising full real-time Feynman path integral molecular dynamics (58) could provide deeper insights into the exact lifetime of these transient tunnelling states.

Second, the structural model is an effective one-dimensional projection of a highly complex three-dimensional biochemical environment. The inverse-physics optimiser successfully deduced the 0.137 nm physical radius of oxygen and the 0.974 nm electrostatic screening length of the confined aAsp/aGlu/aAsn/aHis Grotthuss network. However, by tracking the protons as continuous coordinate tensors subject to pairwise steric and Yukawa forces, the model abstracts away the discrete quantum chemistry of the water wire. Explicit atomistic phenomena such as the discrete formation and cleavage of hydrogen bonds, the transition between Eigen 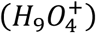 and Zundel 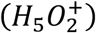 complex geometries and the subsequent rotational reorganisation of water molecules (Bjerrum defects) necessary to reset the wire are mathematically coarse-grained into the effective friction (*γ*) and steric interaction (*A*_*s*_) parameters (22,23).

Finally, the thermodynamic driving force mimicking the biological torque of the *F*_*o*_ rotor was modelled as a continuous, constant external voltage (*F*_*ext*_). In physiological reality, the rotation of the lipid-embedded c-ring against the a-subunit imposes a complex, load-dependent and discrete stepping mechanism (59,60).

However, none of these mathematical simplifications invalidate the core biophysical finding of this study. Introducing explicit 3D hydration dynamics, discrete c-ring stepping or full path-integral coherence would only increase the computational complexity and structural rigidity of the confinement zone. The fundamental proof that classical, non-interacting particles mathematically detonate upon contact with the 1/*r*^12^ steric boundaries of a 1.0 nm Grotthuss wire remains a physical limit. Thus, while future multi-scale models will be required to resolve the explicit atomistic chemistry of the channel, the baseline thermodynamics mathematically dictate that physiological proton flux is sustained exclusively by many-body quantum percolation.

## Data and Code Availability

Data and code will be shared upon reasonable request.

## Author contributions

I.A. Conceptualisation, Data Curation, Formal Analysis, Investigation, Methodology, Software, Validation, Visualisation, Writing – original draft, Writing – review & editing. A.P.L. Formal Analysis, Writing – review & editing.

## Declaration of interests

The authors declare no competing interests.

## Acknowledgements

The authors received no funding for this work. Assistance from Gemini 3.1 (Google) was used to improve the clarity, grammar and conciseness of the manuscript text.

## Supplemental Methods

### Analytical Formulation of the Asymmetric Ratchet Potential

Below is the dimensionless periodic asymmetric ratchet potential U(x) used in the Quantum Langevin Equation. Based on the framework established by Fornes (1), the continuous spatial potential is defined analytically as:

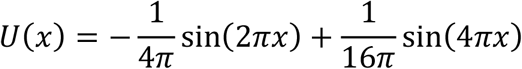

To avoid numerical differentiation artifacts at the steep spatial barriers of the ratchet, the deterministic classical force *F*(*x*) acting on the proton was implemented utilising the exact analytical first derivative:

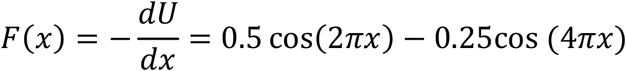

Furthermore, the spatial quantum tunnelling force and thermodynamic diffusion scaling require precise local concavity measurements. The exact second and third analytical derivatives implemented in the numerical integration scheme are:

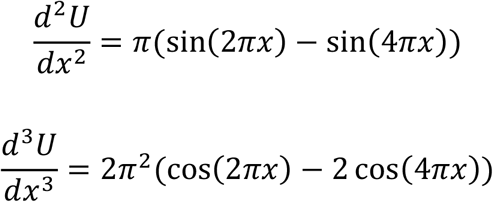

## Notes

### Competing Interest Statement

The authors have declared no competing interest.

## References

1. Mitchell P. Coupling of phosphorylation to electron and hydrogen transfer by a chemiosmotic type of mechanism. Nature. 1961 Jul 8;191:144–8. doi:10.1038/191144a0 PubMed PMID: 13771349.

2. Boyer PD. The ATP synthase--a splendid molecular machine. Annu Rev Biochem. 1997;66:717–49. doi:10.1146/annurev.biochem.66.1.717 PubMed PMID: 9242922.

3. Kühlbrandt W. Structure and Mechanisms of F-Type ATP Synthases. Annu Rev Biochem. 2019 Jun 20;88(Volume 88, 2019):515–49. doi:10.1146/annurev-biochem-013118-110903

4. Junge W, Nelson N. ATP synthase. Annu Rev Biochem. 2015;84:631–57. doi:10.1146/annurev-biochem-060614-034124 PubMed PMID: 25839341.

5. Noji H, Yasuda R, Yoshida M, Kinosita K. Direct observation of the rotation of F1-ATPase. Nature. 1997 Mar 20;386(6622):299–302. doi:10.1038/386299a0 PubMed PMID: 9069291.

6. Yasuda R, Noji H, Kinosita K, Yoshida M. F1-ATPase is a highly efficient molecular motor that rotates with discrete 120 degree steps. Cell. 1998 Jun 26;93(7):1117–24. doi:10.1016/s0092-8674(00)81456-7 PubMed PMID: 9657145.

7. Kinosita K, Yasuda R, Noji H, Adachi K. A rotary molecular motor that can work at near 100% efficiency. Philos Trans R Soc B Biol Sci. 2000 Apr 29;355(1396):473–89. doi:10.1098/rstb.2000.0589 PubMed PMID: 10836501; PubMed Central PMCID: PMC1692765.

8. Diez M, Zimmermann B, Börsch M, König M, Schweinberger E, Steigmiller S, et al. Proton-powered subunit rotation in single membrane-bound F0F1-ATP synthase. Nat Struct Mol Biol. 2004 Feb;11(2):135–41. doi:10.1038/nsmb718 PubMed PMID: 14730350.

9. Yoshida M, Muneyuki E, Hisabori T. ATP synthase--a marvellous rotary engine of the cell. Nat Rev Mol Cell Biol. 2001 Sep;2(9):669–77. doi:10.1038/35089509 PubMed PMID: 11533724.

10. Medvedev ES, Stuchebrukhov AA. Proton diffusion along biological membranes. J Phys Condens Matter. 2011 May;23(23):234103. doi:10.1088/0953-8984/23/23/234103

11. Medvedev ES, Stuchebrukhov AA. Mechanism of long-range proton translocation along biological membranes. FEBS Lett. 2013 Feb 14;587(4):345–9. doi:10.1016/j.febslet.2012.12.010 PubMed PMID: 23268201; PubMed Central PMCID: PMC4222192.

12. Georgievskii Y, Medvedev ES, Stuchebrukhov AA. Proton transport via coupled surface and bulk diffusion. J Chem Phys. 2002 Jan 22;116(4):1692–9. doi:10.1063/1.1428350

13. Nesterov SV, Yaguzhinsky LS, Vasilov RG, Kadantsev VN, Goltsov AN. Contribution of the Collective Excitations to the Coupled Proton and Energy Transport along Mitochondrial Cristae Membrane in Oxidative Phosphorylation System. Entropy. 2022 Dec 13;24(12):1813. doi:10.3390/e24121813 PubMed PMID: 36554218; PubMed Central PMCID: PMC9778164.

14. Heberle J, Riesle J, Thiedemann G, Oesterhelt D, Dencher NA. Proton migration along the membrane surface and retarded surface to bulk transfer. Nature. 1994 Aug;370(6488):379–82. doi:10.1038/370379a0

15. Flegel H, Variyam AR, Amdursky N, Steinem C. ATP synthase activity boosts membrane proton acceptance and lateral diffusion. Proc Natl Acad Sci. 2026 Mar 10;123(10):e2510444123. doi:10.1073/pnas.2510444123

16. Adeniran I, Degens H. Nanoscale spatial confinement of proton flux by cardiolipin drives high-speed lateral proton transport in mitochondria. Manuscript submitted for publication.

17. Feniouk BA, Kozlova MA, Knorre DA, Cherepanov DA, Mulkidjanian AY, Junge W. The Proton-Driven Rotor of ATP Synthase: Ohmic Conductance (10 fS), and Absence of Voltage Gating. Biophys J. 2004 Jun 1;86(6):4094–109. doi:10.1529/biophysj.103.036962

18. Macdonald JE, Ashby PD. The molecular mechanism of ATP synthase constrains the evolutionary landscape of chemiosmosis. Biophys J. 2025 Jul 1;124(13):2103–19. doi:10.1016/j.bpj.2025.05.017

19. Fornés JA. Quantum Ratchets. In: Fornés JA, editor. Principles of Brownian and Molecular Motors [Internet]. Cham: Springer International Publishing; 2021 [cited 2026 Jun 19]. p. 123–48. Available from: https://doi.org/10.1007/978-3-030-64957-9_8 doi:10.1007/978-3-030-64957-9_8

20. Agmon N. The Grotthuss mechanism. Chem Phys Lett. 1995 Oct 13;244(5):456–62. doi:10.1016/0009-2614(95)00905-J

21. Marx D. Proton transfer 200 years after von Grotthuss: insights from ab initio simulations. Chemphyschem Eur J Chem Phys Phys Chem. 2006 Sep 11;7(9):1848–70. doi:10.1002/cphc.200600128 PubMed PMID: 16929553.

22. Köfinger J, Hummer G, Dellago C. Single-file water in nanopores. Phys Chem Chem Phys. 2011 Aug 16;13(34):15403–17. doi:10.1039/C1CP21086F

23. Hassanali A, Giberti F, Cuny J, Kühne TD, Parrinello M. Proton transfer through the water gossamer. Proc Natl Acad Sci. 2013 Aug 20;110(34):13723–8. doi:10.1073/pnas.1306642110

24. Hille B. Ion Channels of Excitable Membranes. 3rd Edition. Sinauer Associates; 2001.

25. Dror RO, Dirks RM, Grossman JP, Xu H, Shaw DE. Biomolecular simulation: a computational microscope for molecular biology. Annu Rev Biophys. 2012;41:429–52. doi:10.1146/annurev-biophys-042910-155245 PubMed PMID: 22577825.

26. Chowdhury D. Stochastic mechano-chemical kinetics of molecular motors: A multidisciplinary enterprise from a physicist’s perspective. Phys Rep. 2013 Aug 1;Stochastic mechano-chemical kinetics of molecular motors: A multidisciplinary enterprise from a physicist’s perspective 529(1):1–197. doi:10.1016/j.physrep.2013.03.005

27. Ankerhold J, Pechukas P, Grabert H. Strong Friction Limit in Quantum Mechanics: The Quantum Smoluchowski Equation. Phys Rev Lett. 2001 Aug 6;87(8):086802. doi:10.1103/PhysRevLett.87.086802

28. Łuczka J, Rudnicki R, Hänggi P. The diffusion in the quantum Smoluchowski equation. Phys Stat Mech Its Appl. 2005 Jun 1;New Horizons in Stochastic Complexity351(1):60–8. doi:10.1016/j.physa.2004.12.007

29. Reimann P. Brownian motors: noisy transport far from equilibrium. Phys Rep. 2002 Apr 1;361(2):57–265. doi:10.1016/S0370-1573(01)00081-3

30. Banerjee D, Bag BC, Banik SK, Ray DS. Solution of quantum Langevin equation: Approximations, theoretical and numerical aspects. J Chem Phys. 2004 May 15;120(19):8960–72. doi:10.1063/1.1711593

31. Machura L, Kostur M, Hänggi P, Talkner P, Luczka J. Consistent description of quantum Brownian motors operating at strong friction. Phys Rev E Stat Nonlin Soft Matter Phys. 2004 Sep;70(3 Pt 1):031107. doi:10.1103/PhysRevE.70.031107 PubMed PMID: 15524506.

32. Caldeira AO, Leggett AJ. Path integral approach to quantum Brownian motion. Phys Stat Mech Its Appl. 1983 Sep 1;121(3):587–616. doi:10.1016/0378-4371(83)90013-4

33. Ford GW, Lewis JT, O’Connell RF. Quantum Langevin equation. Phys Rev A. 1988 Jun 1;37(11):4419–28. doi:10.1103/PhysRevA.37.4419

34. Bezanson J, Edelman A, Karpinski S, Shah VB. Julia: A Fresh Approach to Numerical Computing. SIAM Rev. 2017 Jan;59(1):65–98. doi:10.1137/141000671

35. Shannon RD. Revised effective ionic radii and systematic studies of interatomic distances in halides and chalcogenides. Acta Crystallogr Sect A. 1976;32(5):751–67. doi:10.1107/S0567739476001551

36. Pauling L. The Nature of the Chemical Bond: An Introduction to Modern Structural Chemistry. Ithaca, NY: Cornell University Press; 2010. 664 p.

37. McLaughlin S. The Electrostatic Properties of Membranes. Annu Rev Biophys. 1989 Jun 1;18(Volume 18, 1989):113–36. doi:10.1146/annurev.bb.18.060189.000553

38. Tunuguntla RH, Allen FI, Kim K, Belliveau A, Noy A. Ultrafast proton transport in sub-1-nm diameter carbon nanotube porins. Nat Nanotechnol. 2016 Jul;11(7):639–44. doi:10.1038/nnano.2016.43

39. Swanson JMJ, Maupin CM, Chen H, Petersen MK, Xu J, Wu Y, et al. Proton Solvation and Transport in Aqueous and Biomolecular Systems: Insights from Computer Simulations. J Phys Chem B. 2007 May 1;111(17):4300–14. doi:10.1021/jp070104x

40. Roux B, Schulten K. Computational studies of membrane channels. Structure. 2004 Aug;12(8):1343–51. doi:10.1016/j.str.2004.06.013 PubMed PMID: 15296727.

41. Corry B, Kuyucak S, Chung SH. Tests of continuum theories as models of ion channels. II. Poisson-Nernst-Planck theory versus brownian dynamics. Biophys J. 2000 May;78(5):2364–81. doi:10.1016/S0006-3495(00)76781-6 PubMed PMID: 10777733; PubMed Central PMCID: PMC1300826.

42. Roux B. Statistical mechanical equilibrium theory of selective ion channels. Biophys J. 1999 Jul;77(1):139–53. doi:10.1016/S0006-3495(99)76878-5 PubMed PMID: 10388746; PubMed Central PMCID: PMC1300318.

43. Zwanzig R, Zwanzig R. Nonequilibrium Statistical Mechanics. Oxford, New York: Oxford University Press; 2001. 240 p.

44. Israelachvili JN. Intermolecular and Surface Forces. Academic Press; 2010. 706 p.

45. Purcell EM. Life at low Reynolds number. Am J Phys. 1977 Jan 1;45(1):3–11. doi:10.1119/1.10903

46. Revels J, Lubin M, Papamarkou T. Forward-Mode Automatic Differentiation in Julia [Internet]. arXiv; 2016 [cited 2026 Jun 21]. Available from: http://arxiv.org/abs/1607.07892 doi:10.48550/arXiv.1607.07892

47. Kingma DP, Ba J. Adam: A Method for Stochastic Optimization [Internet]. arXiv; 2017 [cited 2026 Jun 21]. Available from: http://arxiv.org/abs/1412.6980 doi:10.48550/arXiv.1412.6980

48. Glasserman P, Yao DD. Some Guidelines and Guarantees for Common Random Numbers. Manag Sci. 1992 Jun 1;38(6):884–908.

49. Glasserman P. Monte Carlo Methods in Financial Engineering. Springer Science & Business Media; 2013. 603 p.

50. Law A, Kelton WD. Simulation Modeling and Analysis. Boston: McGraw-Hill Education; 2000. 784 p.

51. Kinosita K, Yasuda R, Noji H, Ishiwata S, Yoshida M. F1-ATPase: a rotary motor made of a single molecule. Cell. 1998 Apr 3;93(1):21–4. doi:10.1016/s0092-8674(00)81142-3 PubMed PMID: 9546388.

52. Astumian RD. Thermodynamics and Kinetics of a Brownian Motor. Science. 1997 May 9;276(5314):917–22. doi:10.1126/science.276.5314.917

53. Baradaran R, Berrisford JM, Minhas GS, Sazanov LA. Crystal structure of the entire respiratory complex I. Nature. 2013 Feb 28;494(7438):443–8. doi:10.1038/nature11871 PubMed PMID: 23417064; PubMed Central PMCID: PMC3672946.

54. Wikström M, Krab K, Sharma V. Oxygen Activation and Energy Conservation by Cytochrome c Oxidase. Chem Rev. 2018 Mar 14;118(5):2469–90. doi:10.1021/acs.chemrev.7b00664 PubMed PMID: 29350917; PubMed Central PMCID: PMC6203177.

55. Tunuguntla RH, Allen FI, Kim K, Belliveau A, Noy A. Ultrafast proton transport in sub-1-nm diameter carbon nanotube porins. Nat Nanotechnol. 2016 Jul;11(7):639–44. doi:10.1038/nnano.2016.43 PubMed PMID: 27043198.

56. Yuan Y, Patel RK, Banik S, Reta TB, Bisht RS, Fong DD, et al. Proton Conducting Neuromorphic Materials and Devices. Chem Rev. 2024 Aug 28;124(16):9733–84. doi:10.1021/acs.chemrev.4c00071 PubMed PMID: 39038231.

57. Tegmark M. Importance of quantum decoherence in brain processes. Phys Rev E. 2000 Apr 1;61(4):4194–206. doi:10.1103/PhysRevE.61.4194

58. Markland TE, Ceriotti M. Nuclear quantum effects enter the mainstream. Nat Rev Chem. 2018 Feb 28;2(3):0109. doi:10.1038/s41570-017-0109

59. Ishmukhametov R, Hornung T, Spetzler D, Frasch WD. Direct observation of stepped proteolipid ring rotation in E. coli F0F1-ATP synthase. EMBO J. 2010 Dec 1;29(23):3911–23. doi:10.1038/emboj.2010.259 PubMed PMID: 21037553; PubMed Central PMCID: PMC3020647.

60. Murphy BJ, Klusch N, Langer J, Mills DJ, Yildiz Ö, Kühlbrandt W. Rotary substates of mitochondrial ATP synthase reveal the basis of flexible F1-Fo coupling. Science. 2019 Jun 21;364(6446):eaaw9128. doi:10.1126/science.aaw9128

## References

1. Fornés JA. Quantum Ratchets. In: Fornés JA, editor. Principles of Brownian and Molecular Motors [Internet]. Cham: Springer International Publishing; 2021 [cited 2026 Jun 19]. p. 123–48. Available from: https://doi.org/10.1007/978-3-030-64957-9_8 doi:10.1007/978-3-030-64957-9_8

